# QUALITATIVE ANALYSIS OF CRUCIFEROUS VEGETABLE EXTRACTS USEFUL FOR ESTROGEN METABOLISM FOR DIINDOLYLMETHANE (DIM)

**DOI:** 10.1101/2023.05.09.539959

**Authors:** Joy Ifunanya Odimegwu, Omotuyi Elizabeth Oyinkansola

## Abstract

3,3′-Diindolylmethane (DIM) is a compound derived from the digestion of indole-3-carbinol, found in Cruciferous vegetables (Broccoli, Cabbage, Cauliflower,) which promotes Estrogen metabolism in females. It has been known to help in reduction of heavy blood flow during menstruation especially in females with uterine fibroids. Dim-plus a herbal supplement contains Vitamin E, DIM, Phosphatidlycholine, Spinach, Cabbage, and Broccoli powder. The purpose of this research is to extract and identify the compounds present in Broccoli, Spinach and Cabbage obtained in local markets in Lagos, Nigeria and compare it with the reference standard DIM-plus. Identification of compounds qualitatively by TLC showed different Rf values which were compared with the reference to identify compounds with similar Rf values. Extract was also subjected to HPLC analysis to confirm the presence of DIM in the Cruciferous vegetables using Dim-plus® as standard. Based on the TLC and HPLC analysis it was discovered that the common peak which the crude extracts of the vegetables has is DIM. Thus, the vegetable extracts have Diindolylmethane.

## INTRODUCTION

Herbal medicine which is also called botanical medicine or phytomedicine, refers to the use of plant parts or mixtures of plants extract(s) for medicinal purposes (Russo *et al*., 2013). Examples of such products includes the leaves and roots of *Annona senegalensis*, stem of *Cissus populnea*, the seeds of *Garcinia kola*, the whole plant of *Momordica charantia*, the leaves and roots of both *Newbouldea laevis* and *Rauwolfia vomitoria* (Russo *et al*., 2013).

Medicinal plants are defined as any plants with one or more of its parts containing compounds that can be used for therapeutic purposes or which can be used as precursors for the synthesis of various drugs. Medicinal plants contain numerous biologically active compound such as carbohydrates, proteins, enzymes, fats and oils, minerals, vitamins, alkaloids, quinones, terpenoids, flavonoids, carotenoids, sterols, simple phenolic glycosides, tannins, saponins, polyphenols etc (Sofowora, 1993). Sometimes the living system may require more nutrients than can be received from daily meals and hence the need for dietary supplements.

A dietary supplement is a product that contains a “dietary ingredient” intended to add further nutritional value to (supplement) the diet. It may be one, or any combination of the following substances; a vitamin, a mineral, an herb or other botanical, an amino acid, a dietary substance for use by people to supplement the diet by increasing the total dietary intake, a concentrate, metabolite, constituent, or extract (FDA, 2015). Dietary supplements may be found in many forms such as tablets, capsules, softgels, gelcaps, liquids, or powders. Some dietary supplements can help ensure that you get an adequate dietary intake of essential nutrients; others may help reduce risk of disease (FDA, 2015) and some orders complement the functions of the bodily systems e.g. hormones. A hormone is a naturally occurring substance secreted by specialized cells that affects the metabolism of behaviour of other cells possessing functional receptors for the hormone. A group of hormones, called estrogens, are responsible for the development of female secondary sex characteristics; the development of the breast tissue and the proliferation of the uterine lining. Estrogen helps prepare the body for ovulation. Excess oestrogen is common amongst women in North America (Marcheggiani, 2015).

Stress, caffeine intake, synthetic estrogens in birth control pills and hormone replacement therapy and xeno-estrogens from cleaning products, plastics and cosmetics are among some of the causes of excess levels of estrogen in the body. Due to these environmental factors, many women suffer from “Estrogen Dominance”. (Marcheggiani, 2015, Marquardt, 2019). This manifests as stubborn weight gain, anxiety, premenstrual symptoms of breast tenderness, acne, irritability, fatigue and brain fog. Estrogen dominance can contribute to worsening of health conditions such as infertility, fibrocystic breast, repeated miscarriages, uterine fibroids and endometriosis as well as increase the risk of developing certain cancers. Estrogen detoxification can be achieved effectively through a healthy diet that aims at improving estrogen clearance in the liver and regulation of the action of estrogen at the cellular level. Diets which can be taken includes Cruciferous vegetables; cabbage, cauliflower, broccoli, brussel sprouts, kale, spinach, collard greens and other leafy greens are rich in a nutrient called indole-3-carbinol, or 13C. Indole -3-carbinol gets converted to Di Indolyl Methane (DIM) in the body, which is responsible for clearance of excess estrogens in the liver (Marcheggiani, 2015).

Diindolylmethane is found in many Brassica vegetables through the parent compound glucobrassicin (Aggarwal *et al*., 2007). Ingested glucobrassicin is catalysed via the enzyme myrosinase (stored in vegetables) and turns into indole-3-carbinol which is rapidly digested into both DIM and various other metabolites in the human stomach via acid mediated condensation reactions (Grose and Bjeldanes, 1992). We aim to carry out quality assurance chemical analysis of a known medicinal dietary products/supplement; DIM-plus purported to contain *Brassica oleracea* var. italica, *Brassica oleracea*, and *Basella alba using p*hytochemical analysis including HPLC.

DIM has been implicated in modifying pre-existing estrogen steroids into other metabolites. The process of 2-hydroxylation, likely secondary to aryl hydrocarbon receptor (AhR) activation, may increase the ratio of 2-hydroxyestrone to 16α-hydroxyestrone which is thought to be a less estrogenic profile of estrogen steroids. The processes of 4-hydroxylation and 16-hydroxylation do not appear significantly affected (Okino and Parkin, 2009).

## METHODOLOGY

Plant Materials: *Brassica oleracea var*.*italic* (Broccoli) BOI, *Basella alba* (Indian spinach) BA, *Brassica oleracea* (wild cabbage) BO were purchased fresh from the vegetable garden city, Idiaraba. The species were authenticated at Department of botany, University of Lagos, Akoka, Lagos state, Nigeria. With authentication number: *Brassica oleracea* var. *Italica* BOI (7587), *Brassica oleracea* BO (7589), *Basella alba* BA (7588). The plants were oven dried at 40°C, then the sizes were reduced to powders using industrial grinder. BOI, BO and BA powders were macerated in 80% Methanol and extracted with frequent swirling . The dried extracts were collected, weighed and preserved in a dry bottle until further use.

### THIN LAYER CHROMATOGRAPHY

Preparative TLC was carried out on silica gel coated plates (Merck aluminium sheet silica gel 60F.254), the fractions were separated using the solvent mixture as Hexane: Ethylacetate (80:20 %) After drying the plates, visualization was achieved at UV 254nm and 366nm and spray reagent containing distilled water 8%, Chilled acetone 2%, Perchloric acid 1%, Anisaldehyde 0.5% (8: 2: 1: 0.5) was used. After spraying, heat was applied to dry at about 110 C for 3-5 minutes and the coloured spots obtained were, recorded and the Rf was calculated. The TLC plate was observed under UV-light at 366nm and 254nm wavelengths.

### HPLC ANALYSIS

The HPLC conditions are Agilent 1100. The column used is Zorbax XDB of 5μ. The mobile phase was acetonitrile: water (40 : 60)% using isocratic runs at flow rate of 0.5/minutes. The detector used is UV at wavelength of 254 nm and the column dimension is 150 × 4.6 m.

## Results

**Table 1.**
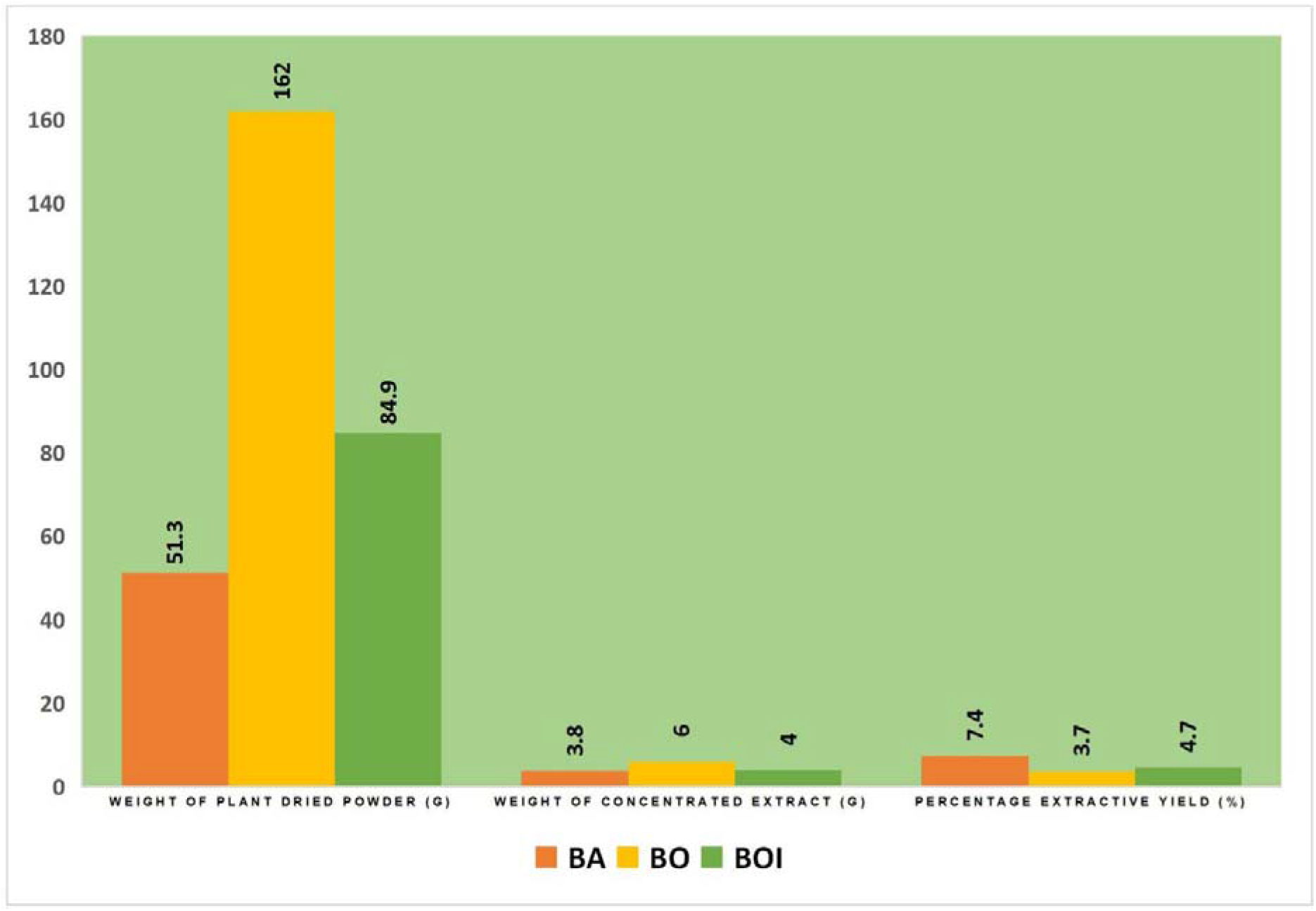
Calculated yields of crude extracts of selected vegetables

## Discussion and conclusion

The HPLC analysis of the extracts of BO, BOI, BA, VE, DP and DI (Fig3-6) showed results of each of the plant extracts from which different peaks at different retention time and the plant with similar compounds were confirmed by comparing the retention times of the peaks with the reference standard chromatograph which is DIM-plus. The methanolic extract of BA detected peak at the retention time of 2.60, 3.51, 7.23, 11.6 minutes. The methanolic extract of BOI detected peaks at the retention time of 2.59, 3.53, 7.25, 11.49 minutes respectively. The n-hexane extract of VE gave peak at retention time of 2.61, 3.50, 7.24 and 11.73 respectively. The methanolic extract of DI detected peaks at the retention time of 2.70, 5.49, 7.24 and 12.22 minutes respectively . The methanolic extract of DIM-plus showed peak at retention time of 2.61, 3.50, 7.24 and 11.73 respectively. From the obtained results, it can be concluded that the qualitative analysis of the methanolic extracts of *Basella alba, Brassica oleracea, Brassica oleracea* var *italica* in a dietary supplement was successful as the bioactive compounds were present in the vegetable extracts.

**Fig. 1.**
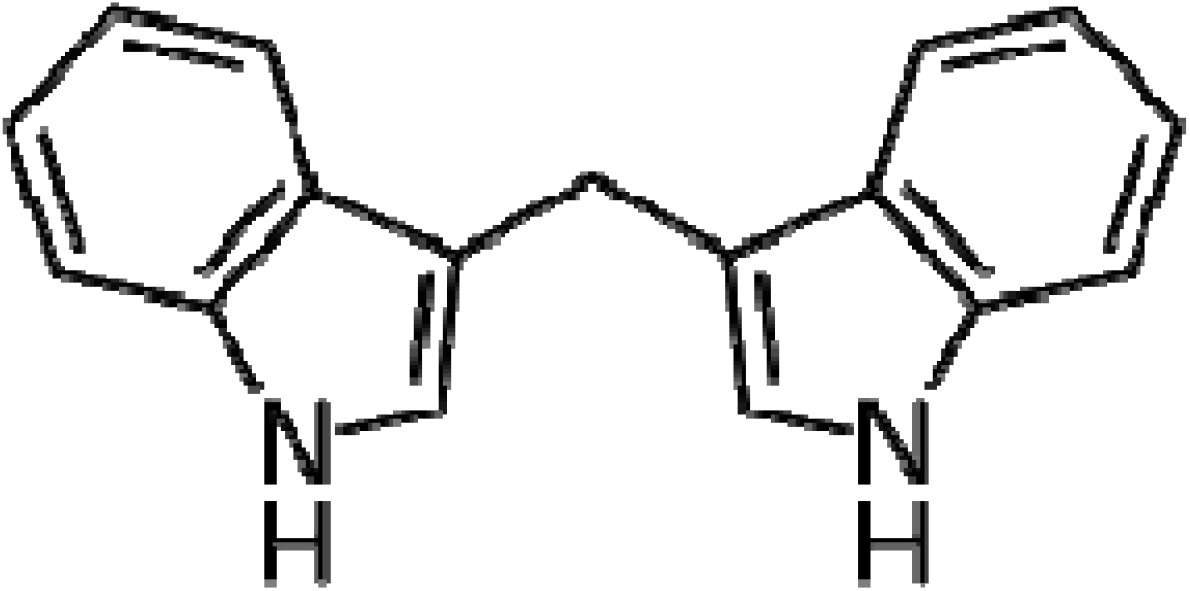
Chemical structure of Diindolylmethane (DIM)

**Fig. 2.**
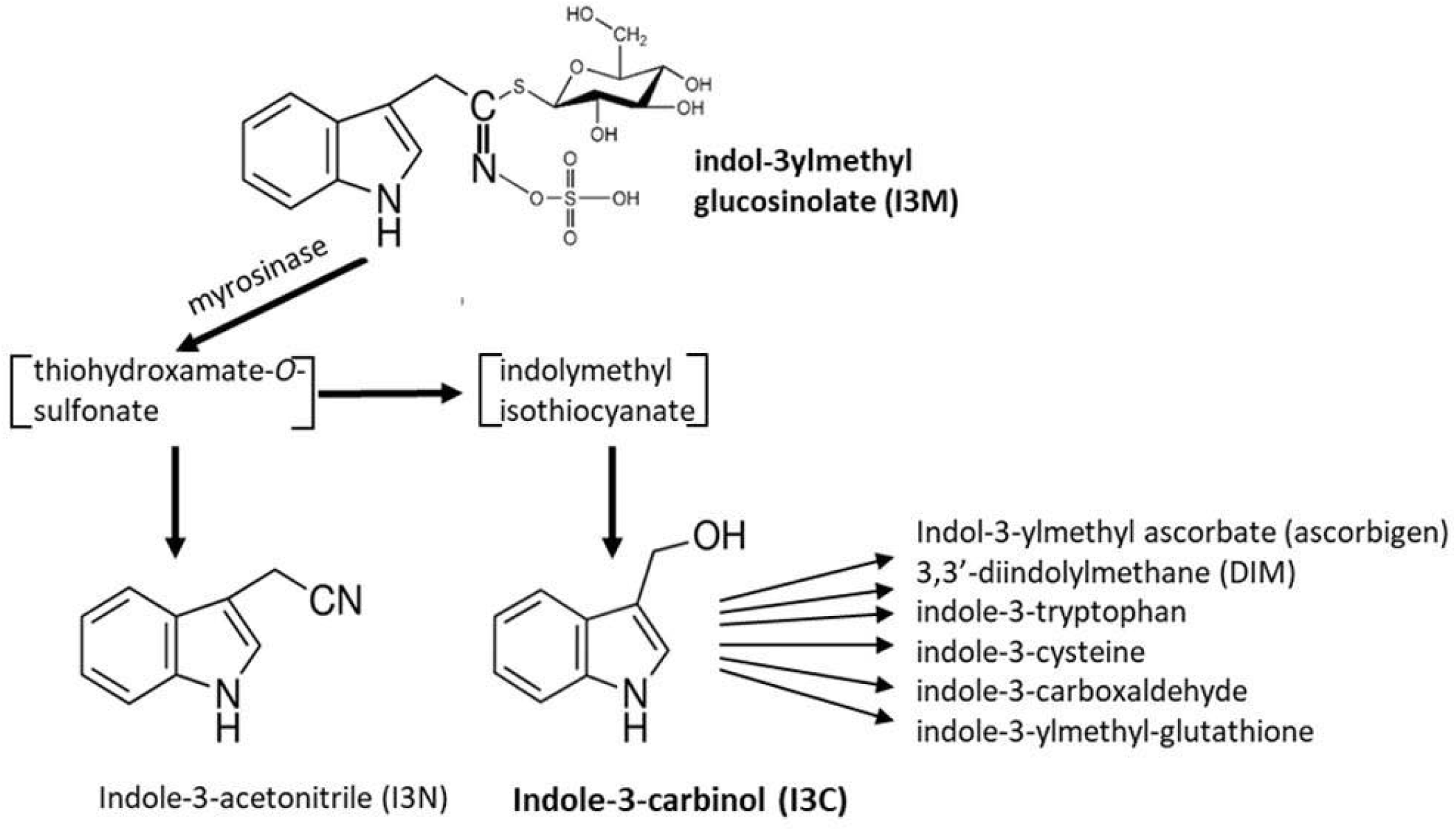
Biosynthesic pathway of Diindolylmethane

**Figure 3.**
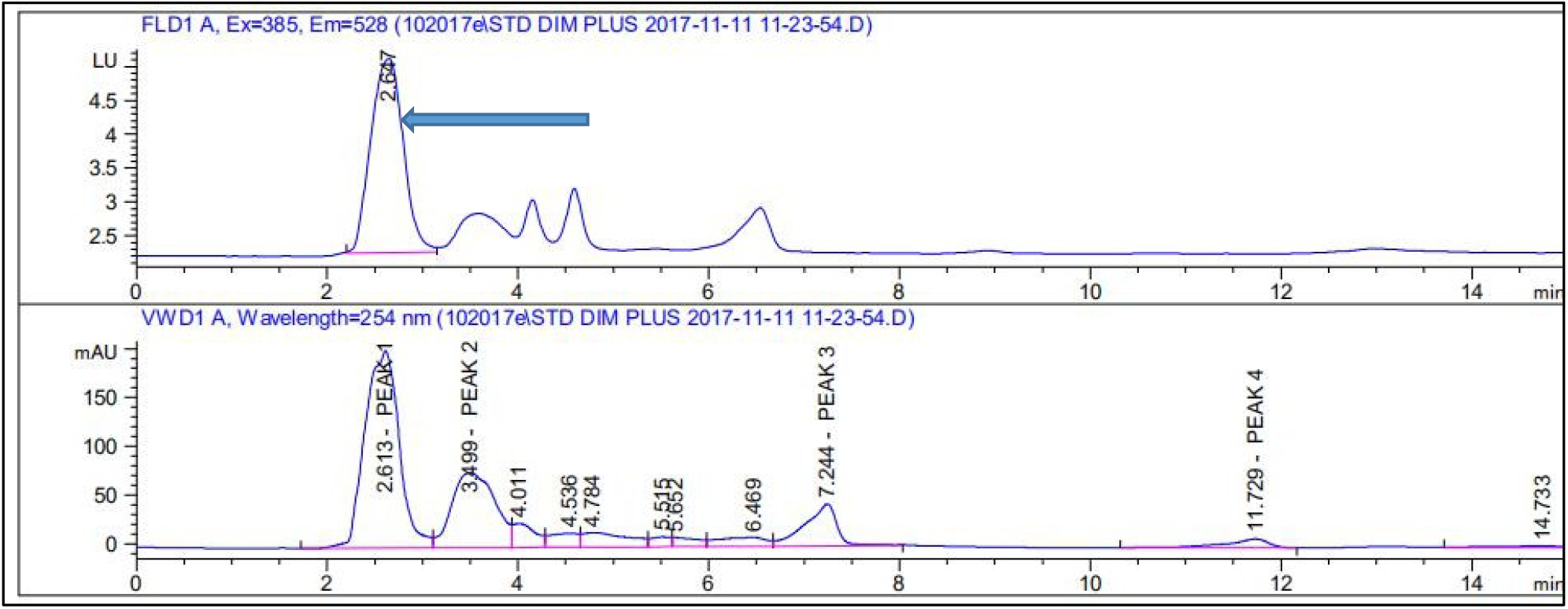
Dimplus caliberation curve obtained from HPLC. *Arrow: DIM peak*

**Figure 4.**
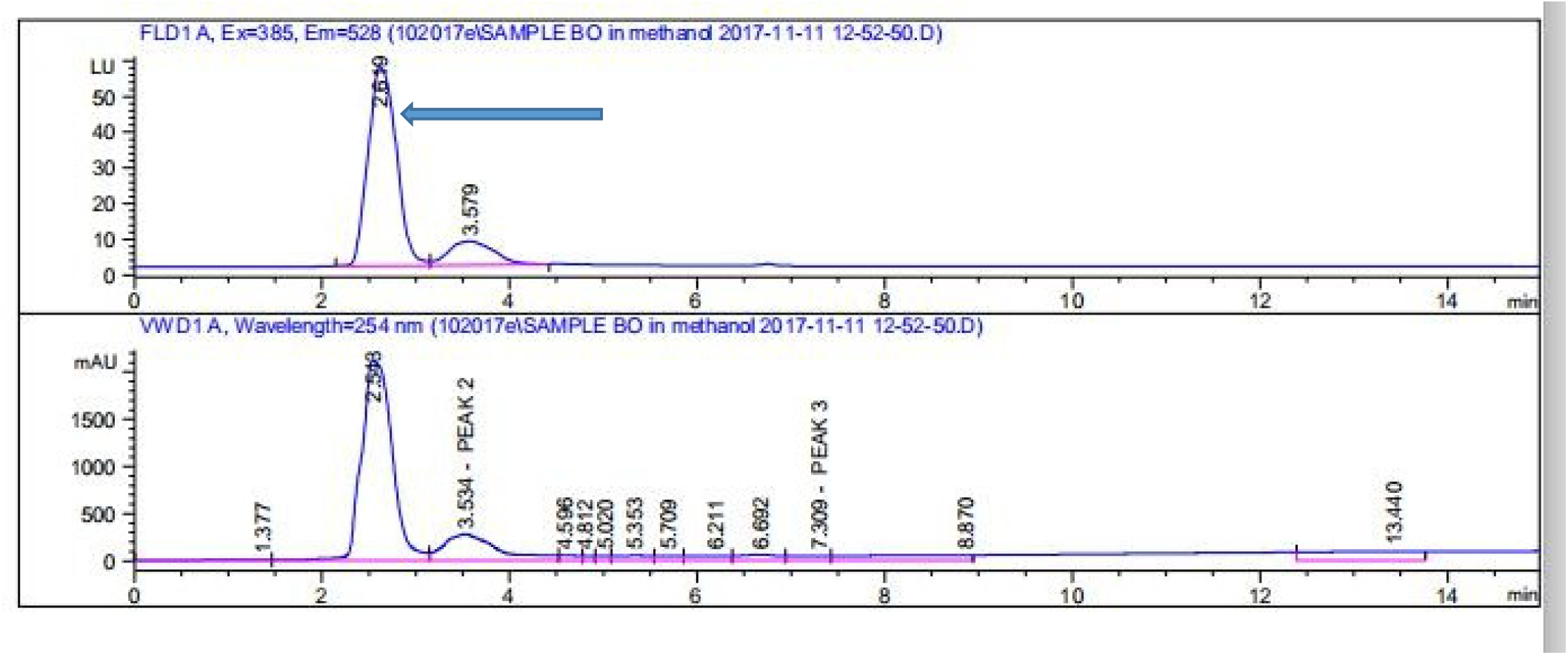
*Brassica oleracea* BO caliberation curve obtained from HPLC. *Arrow: DIM peak*

**Fig 5.**
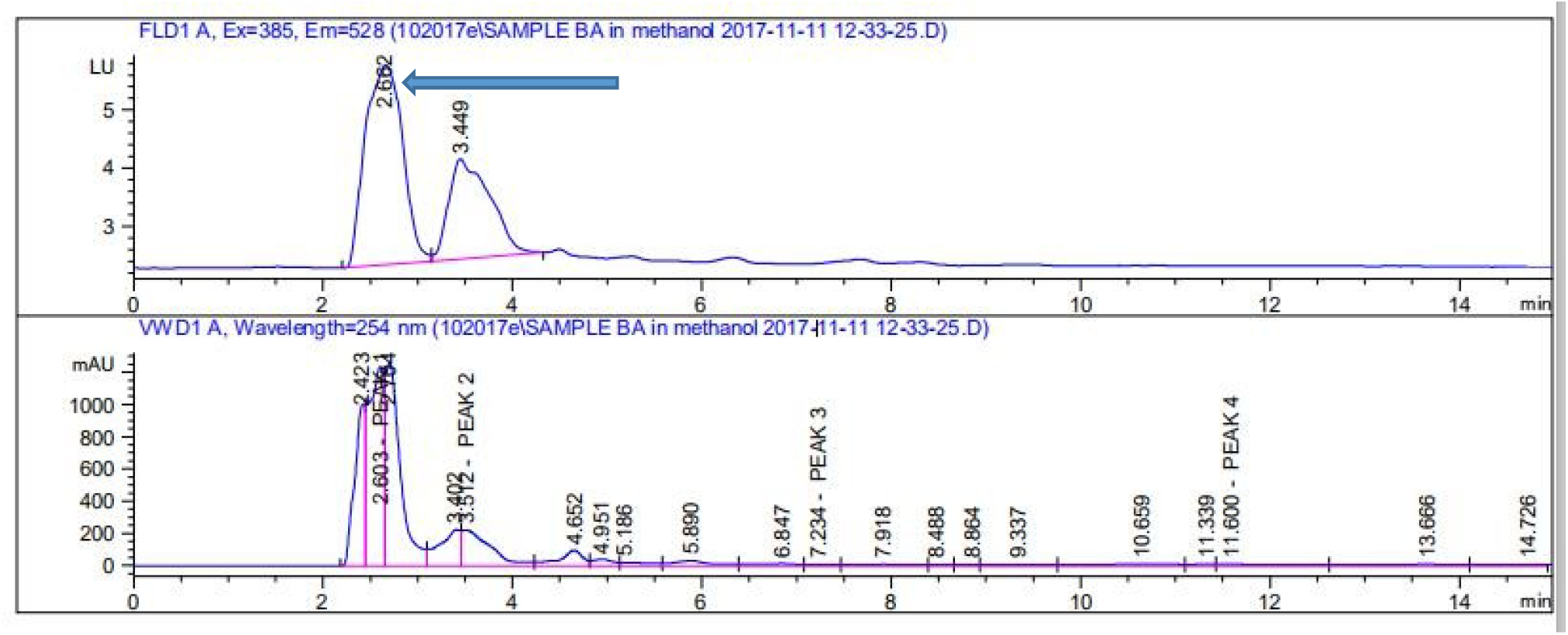
*Basella alba* BA caliberation curve obtained from HPLC. *Arrow: DIM peak*

**Fig. 6.**
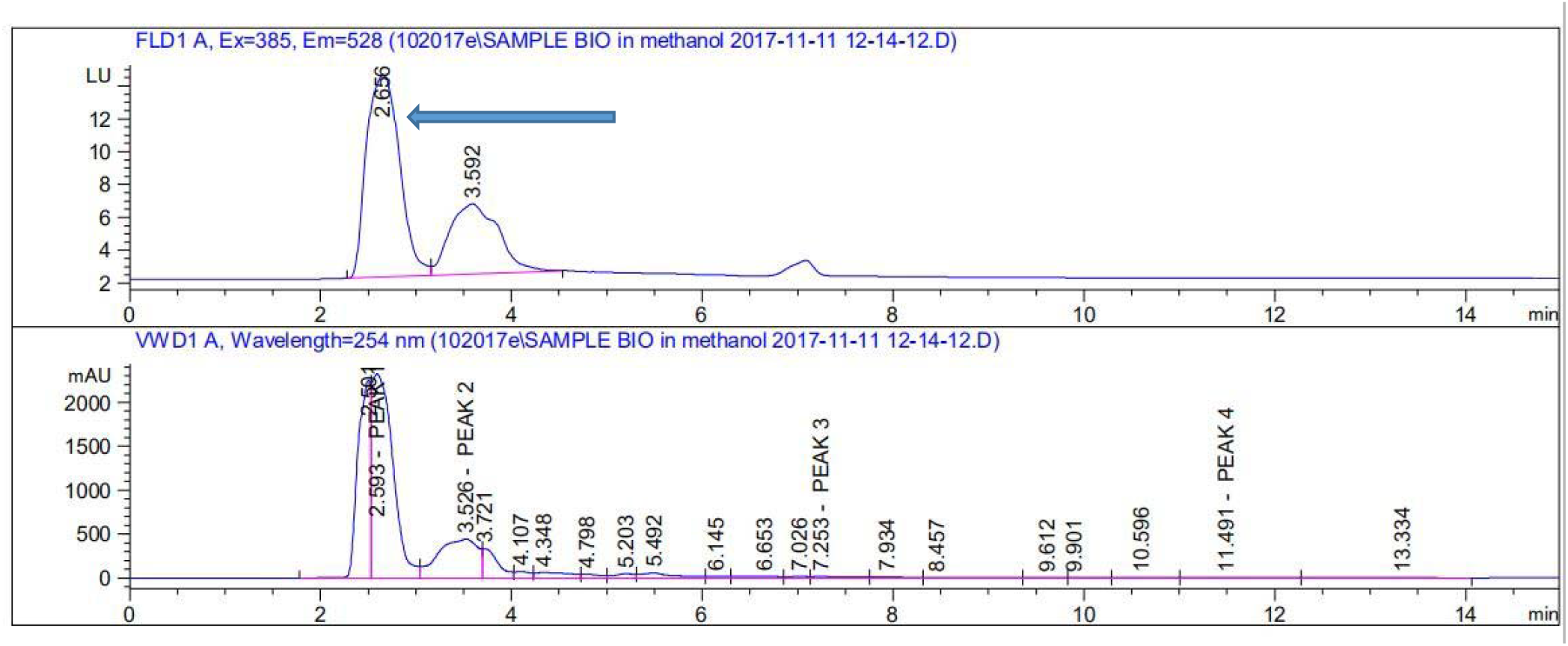
*Brassica oleracea var*.*italic* BIO caliberation curve obtained from HPLC. *Arrow: DIM peak*

